# Loss of zinc transporters ZIP1 and ZIP3 augments platelet reactivity in response to G protein-coupled receptor agonists and accelerates thrombus formation *in vivo*

**DOI:** 10.1101/2021.11.19.469234

**Authors:** A Elgheznawy, P Öftering, M Englert, F Kaiser, C Kusch, U Gbureck, MR Bösl, B Nieswandt, T Vögtle, HM Hermanns

## Abstract

Zinc (Zn^2+^) is considered as an important mediator for thrombosis and haemostasis. However, our understanding of the transport mechanisms that regulate Zn^2+^ homeostasis in platelets is limited. Zn^2+^ transporters, ZIPs and ZnTs, are widely expressed in eukaryotic cells. Using mice globally lacking ZIP1 and ZIP3 (ZIP1/3 DKO), our aim was to explore the potential role of these well-known Zn^2+^ transporters in maintaining platelet Zn^2+^ homeostasis and in the regulation of platelet function. While ICP-MS measurements indicated unaltered overall Zn^2+^ concentrations in platelets of ZIP1/3 DKO mice, we observed a significantly delayed and less efficient Zn^2+^ release upon thrombin-stimulated platelet activation. This resulted in a hyperactive platelet response not only in response to thrombin, but also towards other G protein-coupled receptor (GPCR) agonists. Immunoreceptor tyrosine-based activation (ITAM)-coupled receptor agonist signalling, however, was unaffected. Augmented GPCR responses were accompanied by enhanced Ca^2+^ signalling and PKC activation. Further functional analysis of ZIP1/3 double deficient mice revealed enhanced platelet aggregation, bigger thrombus volume under flow *ex vivo* and faster *in vivo* thrombus formation. The current study thereby identifies ZIP1 and ZIP3 as important regulators for the maintenance of platelet Zn^2+^ homeostasis and function.

## Introduction

Zinc (Zn^2+^) is an essential micronutrient that represents the second most abundant transition element in the body next to iron (1,2). In the human body, plasma contains 0.1% of total Zn^2+^, while the majority of Zn^2+^ content is present intracellularly (3,4). Cellular Zn^2+^ exists either tightly or loosely protein-bound or in a free or non-proteinous ligand-bound status referred to as mobile or labile zinc (3). Zn^2+^ binds to nearly 10% of human proteins to act as a cofactor for maintaining normal cellular signalling (2,5,6).

Zn^2+^ is enriched in platelets and is primarily stored in their α-granules or bound to metallothionein in the cytosol at concentrations up to 30- to 60-fold higher than in the plasma (7–9). Upon platelet activation, Zn^2+^ is released into the circulation and in the microenvironment of the growing thrombus to modulate the activity of several proteins that are involved in platelet aggregation, coagulation and fibrin clot formation (10–13). Consistent with that, blood serum has higher Zn^2+^ concentrations than plasma, indicating that activated platelets release stored Zn^2+^ during clot formation (14). Furthermore, Zn^2+^ deficiency can result in impaired platelet activity and increased bleeding times (15,16). Therefore, Zn^2+^ acts as an important mediator of haemostasis and thrombosis (10, 17–19).

Studies in recent years have suggested that functions of exogenous Zn^2+^ are distinct depending on the concentration. For instance, at low concentrations (30 μM), exogenous Zn^2+^ works as a coactivator via potentiating platelet responses to GPCR and ITAM-coupled receptor agonists to promote platelet activation (20,21). Consistent with that, increased nutritional Zn^2+^ intake increases platelet reactivity in response to suboptimal doses of thrombin, ADP, collagen and adrenaline (22). At intermediate concentrations (0.1-1 mM), exogenous Zn^2+^ can act as platelet agonist to regulate intracellular signalling, leading to granule secretion and PKC/α_IIb_β_3_-dependent aggregation (20,21,23). On the contrary, at high concentrations (3 mM), extracellular Zn^2+^ impaired platelet aggregation, indicating that Zn^2+^ homeostasis needs to be tightly regulated in platelets as uncontrolled increase of labile Zn^2+^ above physiological limits may lead to cell toxicity (17,24). These observations indicate the existence of Zn^2+^ uptake, storage, and release mechanisms to control Zn^2+^ homeostasis in platelets.

Zn^2+^ cannot permeate through the phospholipid bilayers. Therefore, it requires a regulatory machinery to actively control the transport and availability of Zn^2+^ in the cytoplasm, different intracellular organelles, and extracellular space (17,25). Zn^2+^ transporting proteins and a Zn^2+^-buffer system are the main mechanisms to maintain Zn^2+^ homeostasis. Zn^2+^ transporting proteins involve several non-selective cation channels, exchanger and transporters to regulate Zn^2+^ permeability across the cell membrane. Of these channels, transient receptor potential channels (TRPC6 and TRPM7) are considered to be Zn^2+^ permeable (26,27). In addition, the Na^2+^/Ca^2+^ exchanger (NCX) is reported to facilitate Zn^2+^ movement (28). Although TRP channels and NCX are expressed in platelets and megakaryocytes (MKs)(29,30), nothing is known about their role in the regulation Zn^2+^ homeostasis in platelets. Besides these rather unspecific channels and exchanger, Zn^2+^ homeostasis is more specifically regulated by members of *Slc30a* (zinc transporters; ZnTs) and *Slc39a* (Zrt-Irt like proteins; ZIPs) gene families (31,32). ZIP members act as importers to mediate Zn^2+^ influx into the cytosol thereby increasing cytoplasmic Zn^2+^ levels, while ZnT isoforms work as exporters to control Zn^2+^ efflux thereby reducing cytosolic Zn^2+^ levels (33). Transcripts of several ZIP members are detected in substantial amounts in MKs such as ZIP1, ZIP3, ZIP4, ZIP6, ZIP7, ZIP9 and ZIP10 (7). In addition, different isoforms of ZnTs are expressed in MKs and platelets such as ZnT1, ZnT5, ZnT6, ZnT7 and ZnT9 (7,34). Besides Zn^2+^ transportation, also Zn^2+^ storage and buffering system are important for the regulation of Zn^2+^ bioavailability and accumulation in response to the intracellular metabolic alterations. The best characterized stores for free labile Zn^2+^ are the endoplasmic reticulum, Golgi apparatus, vesicles and secretory granules (9,35). However, free mobile Zn^2+^ is sequestered by the increased amount of high affinity Zn^2+^-binding proteins that exist in cytosol and nucleus. Here proteins of the metallothionein family are of particular relevance which can bind up to seven Zn^2+^ atoms and release them most likely due to alterations in the redox status (36). The mechanism by which intracellular Zn^2+^ homeostasis is maintained in terms of platelet activation is unrevealed. Given that Zn^2+^ transporters are the main regulators for Zn^2+^ uptake, storage and release to sustain Zn^2+^ homeostasis, we were interested to explore the role of two plasma membrane localized Zn^2+^ transporters, ZIP1 and ZIP3, in platelets.

## Materials and methods

### Experimental Animals

Double ZIP1/ZIP3 knockout mice (Slc39a1tm1.1Gka,Slc39a3tm1.1Gka/Mmmh) were originally described by Dufner-Beattie *et al*. (37). Cryopreserved spermatozoa were obtained from MMRRC, University of Missouri (RRID: MMRRC_015979-MU), and rederived by in-vitro-fertilization using C57BL/6J isogenic strain background as oocyte donor according to Takeo, T. *et* Nakagata, N. (38). Heterozygous mice were crossed to obtain wild-type and DKO litters which were further bred as homozygous lines. For experiments, mice from the same generations were used. Sex-matched and age-matched male and/or female mice were used for all experiments. Experiments were conducted according to the German animal protection law and in accordance with good animal practice as defined by the Federation of Laboratory Animal Science Associations (FELASA) and the national animal welfare body GV-SOLAS. Animal studies were approved by the district government of Lower Franconia (Bezirksregierung Unterfranken).

### *In vitro* differentiation of bone-marrow megakaryocytes

*In vitro* differentiation of MKs was carried out as described recently (39) using an antibody mixture in combination with magnetic beads for lineage depletion. The antibody negative fraction was cultured in DMEM (Gibco) containing 50 ng/ml TPO and 100 U/ml rHirudin for 72h, before the cells were enriched by straining them with a 20 μm cell strainer prior to experiments.

### RNA isolation and quantitative RT-PCR

Total RNA was isolated from murine MKs using the NucleoSpin RNA kit (Macherey & Nagel) according to the manufacturer’s instructions including a genomic DNA digest. Reverse transcription up to 1μg of RNA was carried out with the High-Capacity cDNA Reverse Transcription Kit (Life Technologies). Quantitative PCR (qPCR) was performed with SYBR Select Master Mix on a ViiA7 (Life Technologies). Gene expression was calculated by the comparative ΔΔCt-method and normalized to the housekeeping gene Rplp0. All primers had melting temperatures of 58-60°C (Primer Express 3.0, Life Technologies).

### Platelet preparation

Murine blood was collected from the retroorbital plexus into 1.5 ml reaction tube containing 300 μl heparin in TBS (20 U/ml, pH 7.3). To prepare platelet-rich plasma (PRP), 200 μl TBS/heparin were added and blood was centrifuged at 800 rpm for 6 min at RT. Then, supernatant and buffy coat were transferred into a new tube containing 300 μl TBS/heparin and centrifuged at 800 rpm for 6 min at RT. To prepare washed platelets, PRP was centrifuged at 2800 rpm for 5 min at RT. Platelet pellet was gently resuspended in 1 ml Ca^2+^-free Tyrode’s buffer containing PGI2 (0.1 μg/ml) and apyrase (0.02 U/ml) and centrifuged at 2800 rpm for 5 min at RT. Finally, the platelet pellet was resuspended in the appropriate volume of Ca^2+^-free Tyrode’s buffer containing apyrase (0.02 U/ml) to reach the required platelet concentration for experiments.

### Measurements of platelet cation levels using ICP-MS

Washed platelets (1×10^6^ cells/μl) were resuspended into sterile 0.9% saline solution without BSA. The cell suspension was centrifuged at 2000 rpm for 2 min to get platelet pellets which were dissolved in 13.8% ultra-pure HNO_3_ and left for 75 minutes at 90°C. The lysed cells were stored at −20°C until measurements of Zn^2+^, Fe^2+^, Ca^2+^ and Mg^2+^ ion levels with ICP-MS (iCAP RQ, ThermoFisher Scientific, Waltham, USA). Before the measurements, the lysates were diluted 1:20 into ultra-pure water to be at final concentration of 0.69% HNO_3_. Five standard solutions were prepared individually for each element containing 10, 1, 0.1, 0.01 and 0.001 mg/l of Zn^2+^/Fe^2+^/Ca^2+^ ions or 100, 10, 1, 0.1 and 0.01 mg/l Mg^2+^ ions (Sigma-Aldrich, Merck KGaA, Darmstadt Germany). The blank containing 0.69% HNO_3_ was subtracted from each standard solution and sample. The internal Rh standard (Sigma-Aldrich, Merck KGaA, Darmstadt Germany) with a concentration of 10 μg/l was automatically added to each sample by the ICP-MS device. The measured ion concentration in mg/l was multiplied with 20, which means that the displayed ion concentration corresponds to the concentration in the cell lysate solution in 13.8% HNO_3_.

### Measurement of free intracellular Zn^2+^ levels in platelets

Washed platelets (5 × 10^5^ cells/μl) were loaded with FluoZin-3 (2 μM) for 30 min at 37°C in the dark. Fluorophore-loaded platelets were washed, centrifuged at 2000 rpm for 2 min and resuspended with Tyrode’s HEPES buffer. 25μl of loaded platelets were diluted with 1ml of Tyrode’s HEPES buffer. Fluorescence intensity was recorded for an initial 50 sec to get basal Zn^2+^ levels in non-loaded platelet (background F0) and for another 50 sec for loaded platelets (untreated F1). Then, thrombin (0.02 U/ml) was added to the loaded platelet suspension and fluorescence intensity was recorded for another 300 sec (treated F2). The measurements were performed on a BD FACSCelesta (Becton Dickinson). Results were analysed by FlowJo Software (TreeStar, USA) to measure geometric mean fluorescence intensity (Geo-MFI). Retained free intracellular Zn^2+^ after stimulation was determined as percent of (F2-F0)/(F1-F0).

### Flow cytometric analysis of platelet integrin activation, degranulation and glycoprotein expression

Measurement of platelet glycoprotein expression has been described recently (39). Briefly, 50 μl of blood were collected in 300 μl heparin in TBS and 700μl Tyrode’s buffer without Ca^2+^ was added. 50 μl of diluted blood were stained for 15 min at RT with saturating amounts of fluorophore-conjugated antibodies and analysed directly after addition of 500 μl PBS. To analyse platelet activation responses, blood samples were washed twice (2,800 rpm, 5 min, RT) in Tyrode’s buffer without Ca^2+^ and finally resuspended in Tyrode’s buffer containing 2 mM Ca^2+^. Platelets were activated with appropriately diluted agonists for 8 min at 37°C followed by 8 min at RT in the presence of saturating amounts of PE-coupled JON/A (4H5, Emfret Analytics) detecting activated α_IIb_β_3_ integrin and FITC-coupled anti-P-selectin (Wug.E9, Emfret Analytics) antibodies in the dark. The reaction was stopped by addition of 500 μl PBS and samples were analysed with a BD FACSCelesta. Platelets were identified by their forward/side scatter (FSC/SSC) characteristics. Obtained data was analysed using FlowJo (TreeStar, Ashland, OR, USA). Mean fluorescence intensities were normalized to background fluorescence of unstained platelets.

### Aggregometry

200 μl of washed platelets in Tyrode’s buffer containing 2 mM Ca^2+^ at a concentration of 2×10^5^ platelets/μl, were transferred into a cuvette. For all measurements with washed platelets, except those with thrombin as agonist, Tyrode’s buffer was supplemented with 100 μg/ml human fibrinogen. Agonists were added as 100-fold concentrates and light transmission was recorded over 10 min with an Apact 4-channel optical aggregation system (APACT, Hamburg, Germany). For calibration of each measurement, Tyrode’s buffer was set as 100% aggregation and washed platelet suspension was set as 0% aggregation, before the agonist was added.

### Adhesion and spreading assay

Ibidi μ-Slide 8 Well chamber was coated with 100 μg/ml human fibrinogen at 4°C overnight and blocked for at least 1h at RT with 1% BSA in sterile PBS. The wells were rinsed with Tyrode’s buffer and 300 μl washed platelets (250,000 cells/μl in Tyrode’s containing 2 mM Ca^2+^) were stimulated with thrombin (0.01 U/ml) and immediately added to the fibrinogen surface. Platelets were allowed to adhere and spread at 37 °C for 10 min. The non-adhered platelets were washed away and the adhered cells were fixed by addition of 300 μl 4% PFA/PBS for 10 min at RT. Then, the wells were washed twice with 300 μl PBS. Platelets were visualized by differential interference contrast (DIC) microscopy with a Zeiss Axiovert 200 inverted microscope (100x/1.4 oil objective). Representative images were taken using a CoolSNAP-EZ camera (Visitron, Munich, Germany) and evaluated according to different platelet spreading stages with ImageJ (National Institutes of Health, Bethesda, MD, USA). Spreading stages were defined as follows: 1: round; no filopodia, no lamellipodia. 2: only filopodia. 3: lamellipodia; full spreading

### Collagen flow chamber assay

Coverslips (24×60 mm) were coated with 200 μg/ml fibrillar type-I collagen (Horm) overnight at 37°C and blocked for 1 h with 1% BSA at RT. Blood (700 μl) was collected into 300 μl heparin in TBS (20 U/ml, pH 7.3) and two parts of blood were diluted with one part Tyrode’s buffer with 2 mM Ca^2+^. Platelets were labelled with a DyLight-488 conjugated anti-GPIX Ig derivative (0.2 μg/ml; Emfret Analytics) for 5 min at 37°C. The diluted blood was filled into a 1 ml syringe and connected to a transparent flow chamber with a slit depth of 50 μm, equipped with the coated coverslips. Perfusion was performed using a pulse-free pump under high shear stress equivalent to a wall shear rate of 1,000 s-1 for 4 min. Thereafter, coverslips were washed by a 4 min perfusion with Tyrode’s buffer at the same shear rate and phase-contrast and fluorescent images were recorded from at least five different fields of view (63x objective) using a Leica DMI6000 microscope. Image analysis was performed using Image J (NIH). Thrombus formation was expressed as the mean percentage of the total area covered by thrombi (= surface coverage) and as the mean integrated fluorescence intensity (= thrombus volume).

### Measurements of intracellular Ca^2+^ levels

Washed platelets were adjusted to a concentration of approximately 2.5×10^5^/μl in Ca^2+^-free Tyrode’s buffer. 100 μl of this suspension were loaded with 3 μM fura-2-AM F-127 for 25 min at 37°C. After labelling, the platelets were washed once and resuspended in 500 μl Tyrode’s buffer containing 1 mM Ca^2+^ for measurement of Ca^2+^ influx. This cell suspension was transferred into a cuvette and fluorescence was measured with a PerkinElmer LS 55 fluorimeter (Perkin Elmer, Waltham, MA, USA) under stirring conditions. Excitation was switched between 340 and 380 nm and emission was measured at 509 nm. Basal Ca^2+^ levels were recorded for 50 s before the indicated agonist was added. Each measurement was calibrated using 1% Triton X-100 and EGTA.

### Intravital microscopy of FeCl_3_-injured mesenteric arterioles

Mice (16 - 19 g body weight) were anaesthetized and the mesentery was exteriorized through a midline abdominal incision. Arterioles free of fat tissues with a diameter of 35 - 60 μm were visualized using a Zeiss Axiovert 200 inverted microscope (10x/0.25 air objective) equipped with a 100 W HBO fluorescent lamp and a CoolSNAP-EZ camera. Injury was induced by topical application of a 3 mm^2^ filter paper saturated with 20% FeCl_3_. Adhesion and aggregation of fluorescently labelled platelets (achieved by previous intravenous injection of a DyLight-488 conjugated anti-GPIX Ig derivative; Emfret Analytics) in arterioles was monitored for 40 min or until complete occlusion occurred (blood flow stopped for >1 min). Digital images were recorded and analysed using MetaMorph software.

### Western blotting

Washed platelets were adjusted at concentration of 4×10^5^/μl in BSA- and Ca^2+-^free Tyrode’s buffer and supplemented with apyrase (2 mM f.c.). Under stirring conditions in an aggregometer at 37°C, platelets were stimulated with different agonists as indicated in the figures. The stimulation was stopped by mixing the platelet suspension with 10x concentrated ice-cold cell lysis buffer (1x f.c., Cell Signaling Technology #9803) containing Pefabloc SC-Protease-Inhibitor and phosphatase inhibitors (Carl Roth) and kept on ice for at least 30 min. Afterwards, platelet lysates were mixed with 4x SDS sample buffer (1x f.c.), boiled for 5 minutes and subjected to Western blot analysis. Samples were separated by SDS-PAGE using the 10% gel system for electrophoresis, followed by transfer onto a PVDF membrane. Membranes were blocked for 10 minutes in 10% BSA in TBS-T and then incubated overnight at 4°C with the indicated antibodies obtained from Cell Signaling Technology (phospho-PKC Substrate Motif (#6967), phospho-p42/44 MAPK (#4370), total p42/44 (#9102), β-actin (#3700)). Afterwards, the membrane was washed three times with TBS-T at RT. Next, the membranes were incubated with the appropriate secondary HRP-labelled antibodies (CST #7074 or #7076) for 1 h at RT. Finally, the membranes were washed several times and proteins were visualized using the Clarity™ Western ECL Substrate (Bio-Rad) and the ChemiDoc Imaging System (Bio-Rad).

### Statistical analysis

The shown data are expressed as mean ± SEM. When applicable, a modified Student’s t-test was used to analyse differences between unpaired two group’s data. For analysis of more than two groups, one-way Anova or two-way Anova for repeated measures followed by the Bonferroni multiple comparison post-hoc test was applied. p-values <0.05 were considered as statistically significant (*), p<0.01 (**) and p<0.001 (***).

## Results

### Zn^2+^ homeostasis is disturbed in platelet lacking ZIP1 and ZIP3 transporters

Several studies indicated the essential role of Zn^2+^ in the regulation of platelet activity and signalling (19,20). However, the transport mechanisms that control intracellular levels of Zn^2+^ in platelets and their relevance for platelet function remained unaddressed. Our previous study indicated that several members of ZIP (*Slc39a*) family of transporters, which regulate the import of Zn^2+^ into the cytosol (25), are expressed in MKs (7). RNA analysis indicated that ZIP1 and ZIP3 were among the most highly expressed transporters in MKs during the differentiation and maturation stages (7). Therefore, we used a previously described mouse model globally lacking the expression of ZIP1 and ZIP3 transporters (ZIP1/3 DKO) (37). Quantitative RT-PCR analyses clearly demonstrated that *in vitro* differentiated MKs of the double knock-out mice are deficient for the expression of *Slc39a1* and *Slc39a3* (genes encoding ZIP1 and ZIP3, respectively) (**Fig. 1A**). ZIP1/3 DKO mice showed unaltered blood cell (WBCs, RBCs) and platelet counts as compared to wild-type (WT) animals. However, there was a slight but significant reduction in platelet size in ZIP1/3 DKO mice (**Suppl Fig. 1**).

**Figure 1:**
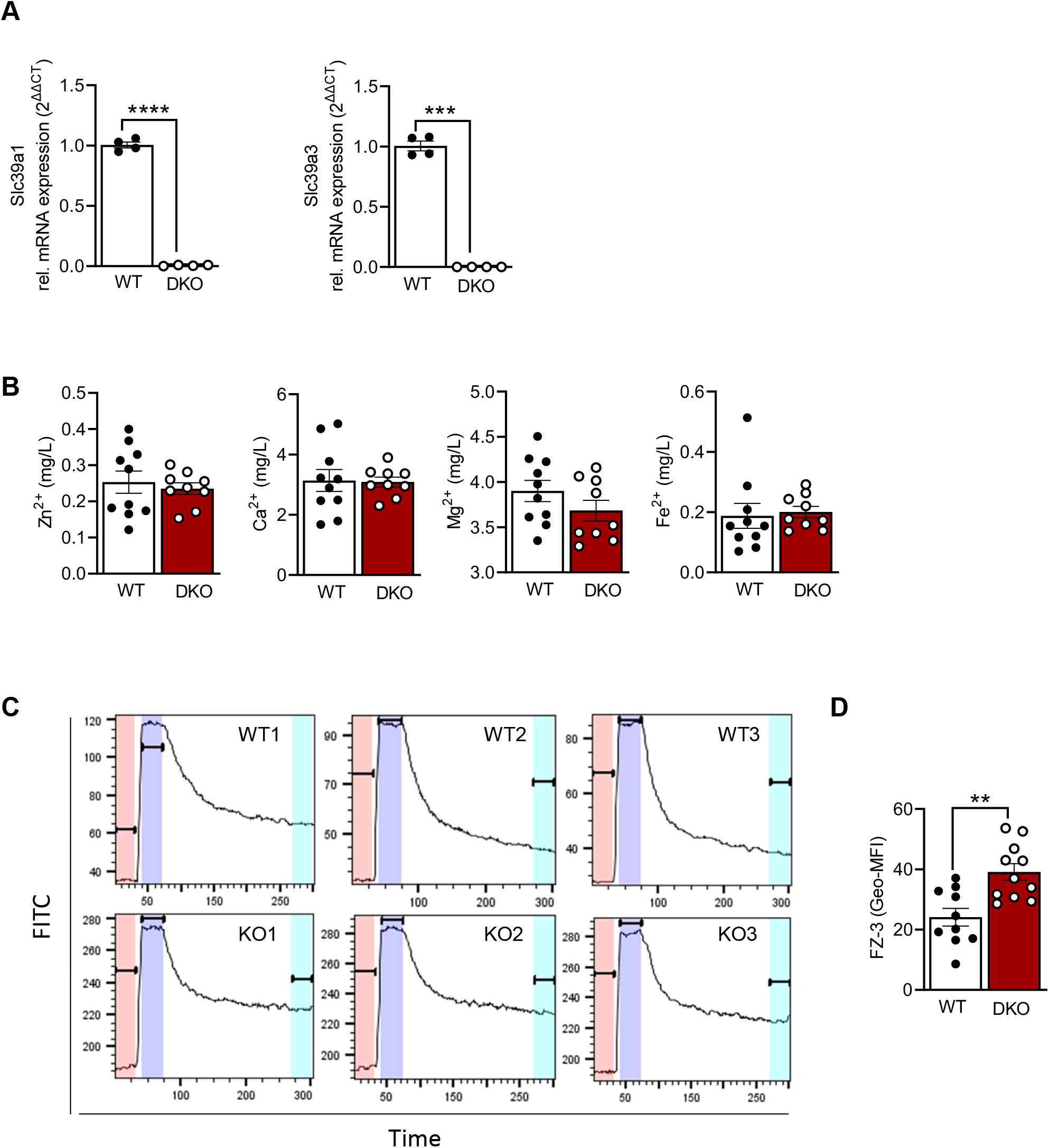
Platelets from ZIP1/3 DKO mice display a delayed and reduced release of free intracellular Zn^2+^ in response to thrombin. (A) Quantitative RT-PCR analysis of mRNA levels of *Slc39a1* and *Slc39a3* in megakaryocytes from wild-type (WT) and ZIP1/3 knockout mice (ZIP1/3 DKO). (B) Determination of intracellular amounts of Zn^2+^, Ca^2+^, Mg^2+^ and Fe^2+^ by ICP-MS analysis of washed platelets. (C) Retained intracellular levels of free Zn^2+^ determined by FluoZin3 fluorescence upon stimulation of WT or ZIP1/3 DKO platelets with thrombin (0.02 U/ml, 5min). n= 10-11, (Students t-test), ****P<0.0001 (D) Representative graphs of FluoZin3 labelling of WT and ZIP1/3 DKO platelets before and after stimulation with thrombin (0.02 U/ml).

Since ZIP1 and ZIP3 act as importers to promote Zn^2+^ influx into the cytosol, we determined total levels of intracellular Zn^2+^ (protein-bound and labile) by inductively coupled plasma mass spectrometry (ICP-MS). No significant differences were found for overall [Zn^2+^]_i_ or other divalent cations ([Ca^2+^]_i_, [Mg^2+^]_i_, [Fe^2+^]_i_) (**Fig. 1B**). To determine the relative ratio of free intracellular Zn^2+^, WT and ZIP1/3 DKO washed platelets were incubated with the Zn^2+^-specific cell permeable fluorescent dye Fluo-Zin3. ZIP1/3 DKO mice express low levels of GFP from the *Slc39a1* and *Slc39a3* gene locus (37) which makes it difficult to compare the basal amount of free [Zn^2+^]_i_ in WT vs DKO mice. The lack of other commercially available specific Zn^2+^ fluorophores, however, necessitated its usage. Despite the slight increase in background fluorescence, it is apparent that platelets from DKO mice contain significant amounts of FluoZin3-stainable [Zn^2+^]_i_ (**Fig. 1C**), which interestingly, is not as efficiently released from the cells in response to thrombin (0.02 U/ml) as in WT platelets (**Fig. 1D**). While WT platelets retained only 24.12% ± 2.942 (n=10) of the initial background corrected FluoZin3 fluorescence, ZIP1/3 DKO platelets retained 39.12% ± 2.766 (n=11; p<0.01).

### Platelets lacking ZIP1 and ZIP3 transporters are hyperreactive to GPCR agonists

Activated platelets release Zn^2+^ into the circulation to modulate proteins that play a major role in platelet adhesion and aggregation (10–13). Previous studies have indicated that Zn^2+^ can act as agonist or second messenger to modulate platelet reactivity in a concentration-dependent manner (19,20). Taken in consideration that disturbed [Zn^2+^]_i_ alters platelet function, we evaluated the aggregation phenotype of platelets lacking ZIP1 and ZIP3 transporters. Interestingly, ZIP1/3 DKO mice displayed an enhanced platelet aggregation in response to threshold concentrations of GPCR agonists such as thrombin (0.003 U/ml), ADP (0.5 μM) and U46619 (0.2 μM) (**Fig.2A**). However, in response to collagen or collagen-related peptide (CRP) (**Fig.2B**), agonists of the ITAM-coupled receptor glycoprotein (GP)VI, there was no difference in platelet aggregation between ZIP1/3 DKO and their corresponding WT controls. This suggested that loss of ZIP1/3 might selectively alter GPCR-coupled signalling in activated platelets.

**Figure 2:**
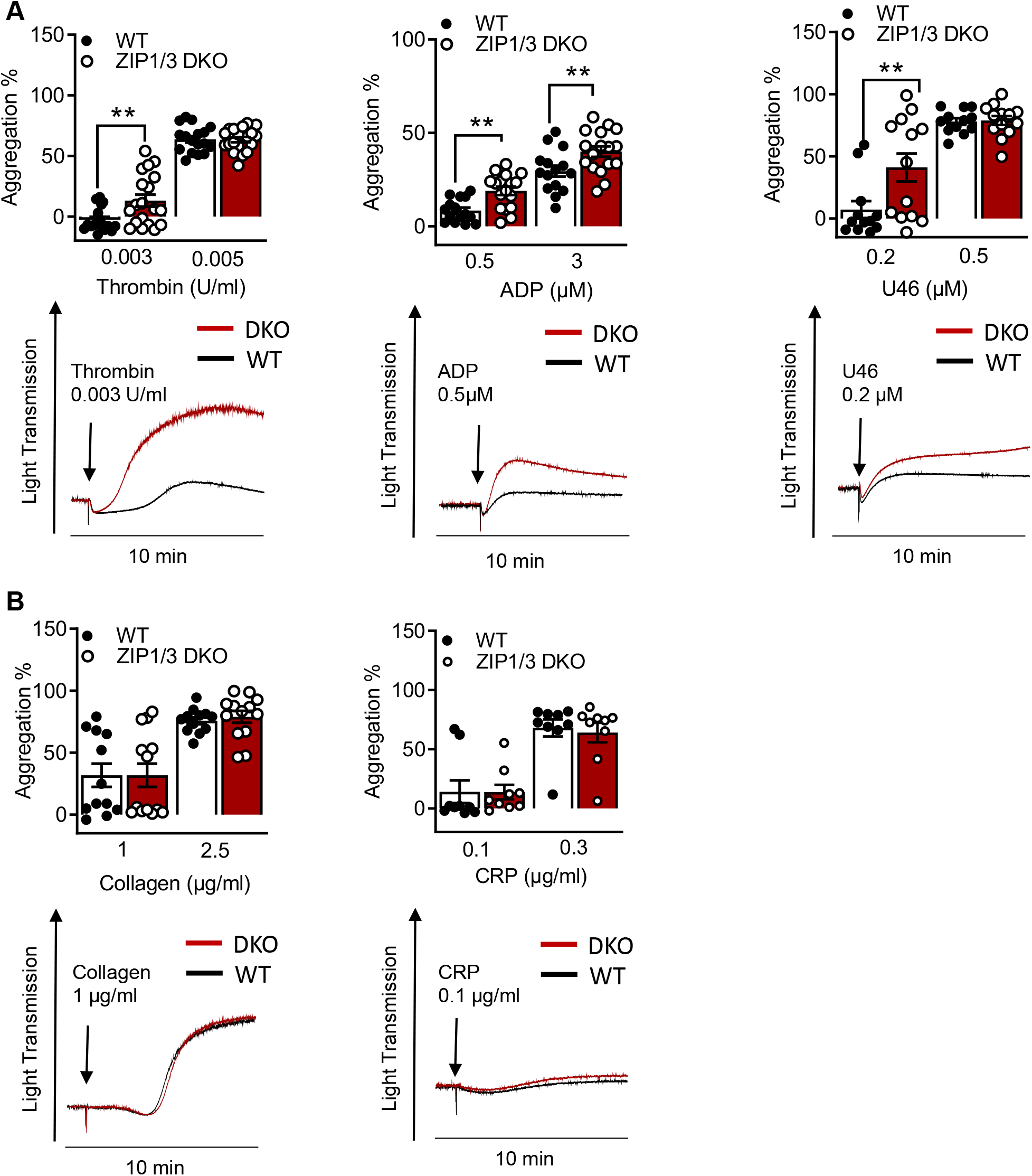
Platelets from ZIP1/3 DKO mice are hyperresponsive towards GPCR agonists but not to ITAM-coupled receptor agonists. Platelet aggregation was measured using Fibrintimer 4-channel aggregometer after the stimulation of washed platelets isolated from WT and ZIP1/3 DKO mice with (A) different GPCR agonists (thrombin, ADP, U46619) and (B) ITAM-coupled receptor agonists (collagen, CRP) using the indicated concentrations; n=9-19 (two-way ANOVA + Bonferroni), **P<0.01.

To corroborate this hypothesis, we measured activation of α_IIb_β_3_ integrin using the JON-A/PE antibody, which only binds α_IIb_β_3_ in its active conformation, in flow cytometry (**Fig. 3A-C**). Our analysis showed that ZIP1/3 deficiency promoted α_IIb_β_3_ integrin activation in response to threshold concentrations of thrombin (0.002 U/ml) (**Fig.3A**), however, responses to ADP, the thromboxane A2 (TxA2) analogue U46619 (**Fig. 3B**) or the ITAM receptor agonist CRP (**Fig.3C**) were unaffected. Similar observations were made for P-selectin exposure (**Fig. 3D-F**). These results indicated that ZIP1 and ZIP3 transporters might be more involved in the regulation of PAR signalling pathway to mediate platelet aggregation. Of note, there was no difference in the total expression levels of integrins (α_IIb_β_3_, α5 and α5) in platelets from ZIP1/3 DKO and WT mice (**Suppl Fig.2A**). Similarly, no difference was detected in the surface expression levels of the GPIb-V-IX complex (**Suppl Fig. 2B**), CD9, CLEC2 and GPVI (**Suppl Fig. 2C**).

**Figure 3:**
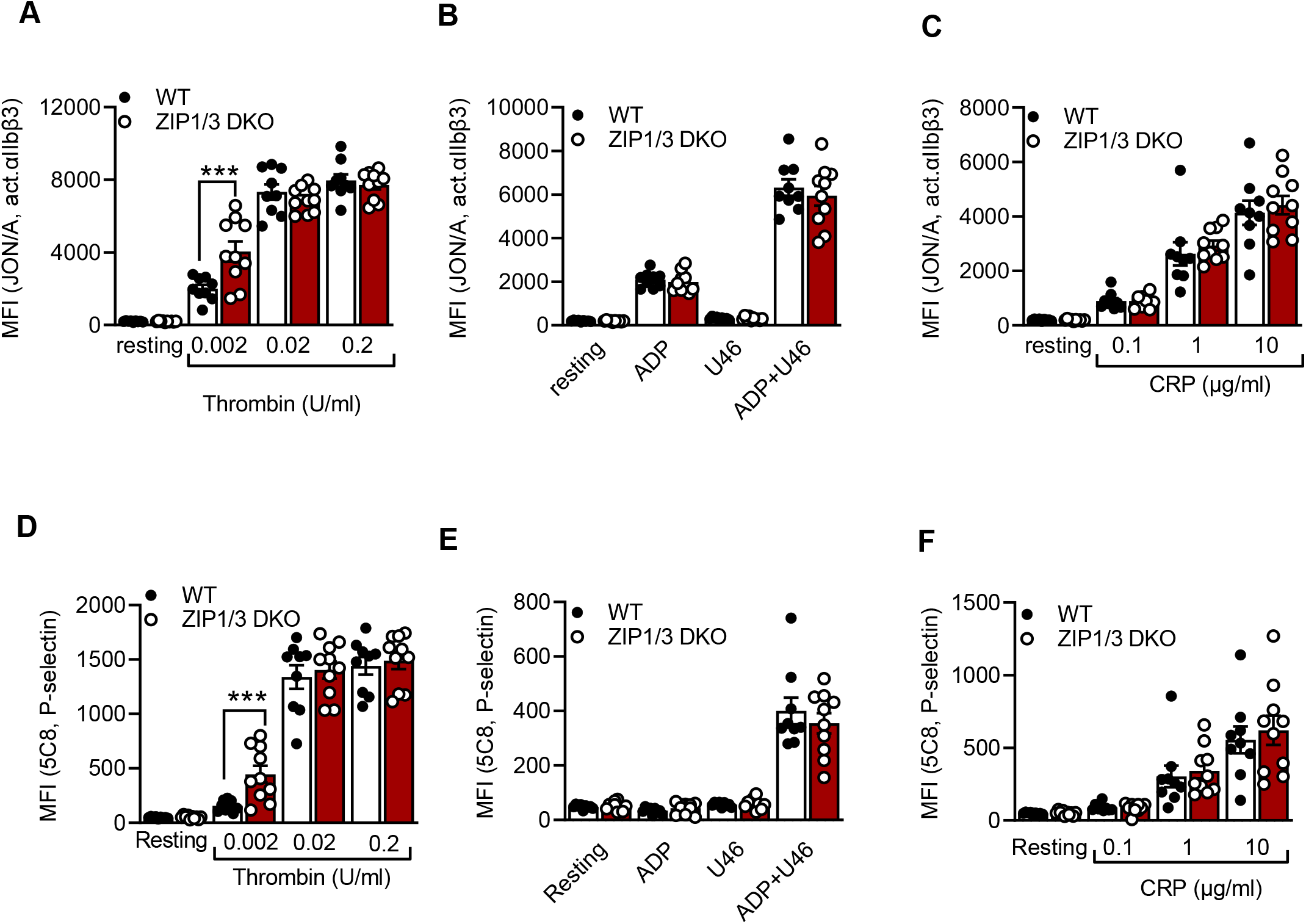
Analysis of platelet integrin activation and degranulation by flow cytometry. Platelets from WT and ZIP1/3 DKO mice were activated with either thrombin (A, D), ADP (10μM), U46619 (3μM) and their combination (B,E) or CRP (D, F) and stained with saturating amounts of PE-coupled JON/A detecting activated αIIbβ3 integrin (A-C) or FITC-coupled anti-P-selectin (D-F). Mean background fluorescence of unstained platelets was subtracted from all data sets. n=9-10 (two-way ANOVA + Bonferroni), ***P<0.001

Since it has been reported that intracellular Zn^2+^ concentrations regulate platelet shape changes (19), we tested platelet adhesion and spreading on fibrinogen in the presence of thrombin under static conditions. However, platelets isolated from ZIP1/3 DKO mice did not show any difference in adhesion and spreading compared to WT mice (**Fig.4A, B**).

**Figure 4:**
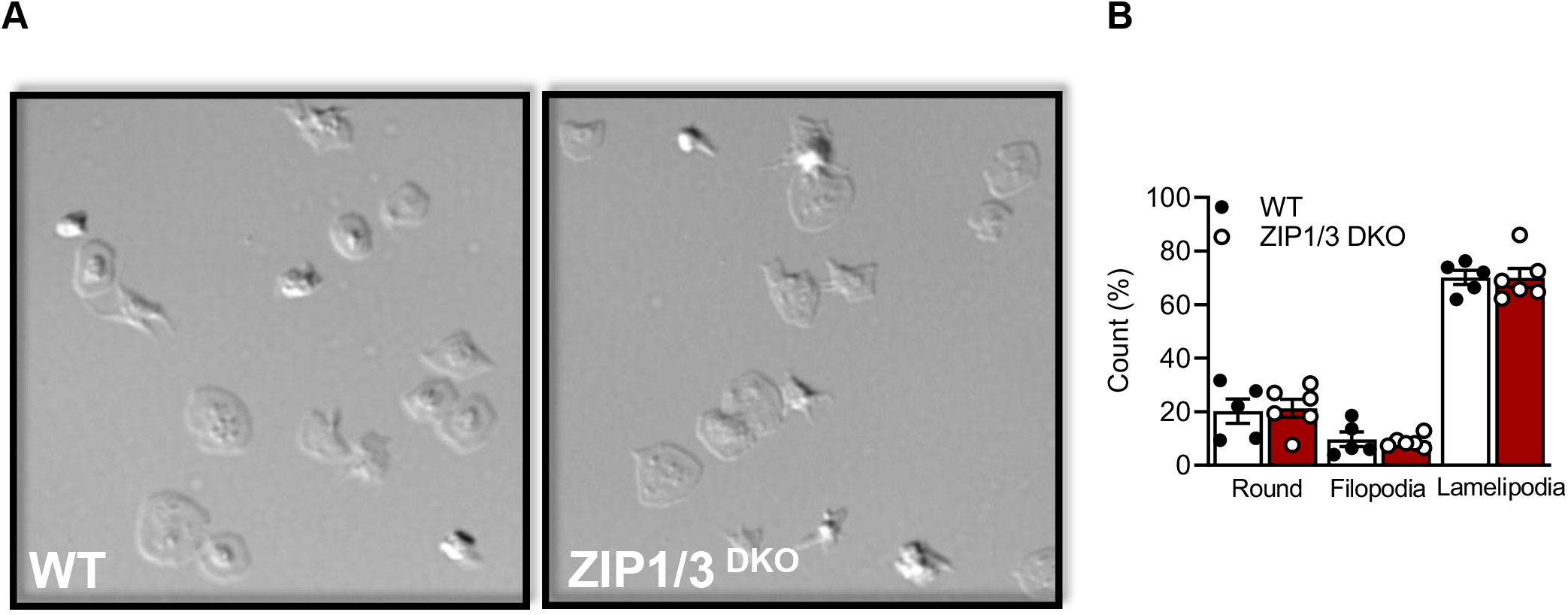
ZIP1/3 deficiency has no effect on *in vitro* platelet adhesion and spreading under static conditions. (A) Washed platelets from WT and ZIP1/3 DKO mice were allowed to adhere on fibrinogen for 10 min in the presence of 0.01 U/ml thrombin. (B) Quantification of spreading stages (round=no filopodia, no lamellipodia)

### ZIP1/3 deficiency increases Ca^2+^ influx and PKC activation in platelets

It has been speculated that Ca^2+^ and Zn^2+^ may work synergistically to induce platelet activation (20). Even though overall [Ca^2+^]_i_ appeared to be equivalent in WT and ZIP1/3 DKO platelets at resting conditions in the absence of extracellular Ca^2+^ as determined by ICP-MS (**Fig. 1B**), Ca^2+^ influx was significantly enhanced in ZIP1/3 DKO mice in response to thrombin (0.01 U/ml) (**Fig. 5A**), while Ca^2+^ influx in response to CRP was unaffected (**Fig. 5B**). Given the increase in Ca^2+^ influx in response to thrombin (**Fig. 5A**) and the fact that increased [Zn^2+^]_i_ has been shown to modulate protein kinase C (PKC) (20), we assessed the importance of platelet ZIP/ZIP3 in agonist-mediated activation of PKC by determination of the phosphorylation status of PKC substrates using Western blot analyses. Deletion of ZIP1/3 resulted in elevated phosphorylation levels of PKC substrates in response to 0.003 U/ml thrombin (**Fig. 6A**), 0.5 μM ADP (**Fig. 6B**) and to a lesser extent to 0.2μM U46619 (**Fig. 6C**) in stimulated platelets compared to platelets from WT mice. As observed in other assays before, the ITAM receptor agonists CRP (0.1μg/ml) (**Fig. 6D**) or collagen (1μg/ml) (**Fig. 6E**) did not induce a stronger phosphorylation of PKC substrates in platelets from ZIP1/3 DKO compared to WT platelets. Therefore, the presence of ZIP1 and ZIP3 transporters appears to increase GPCR’s sensitivity to their respective agonists.

**Figure 5:**
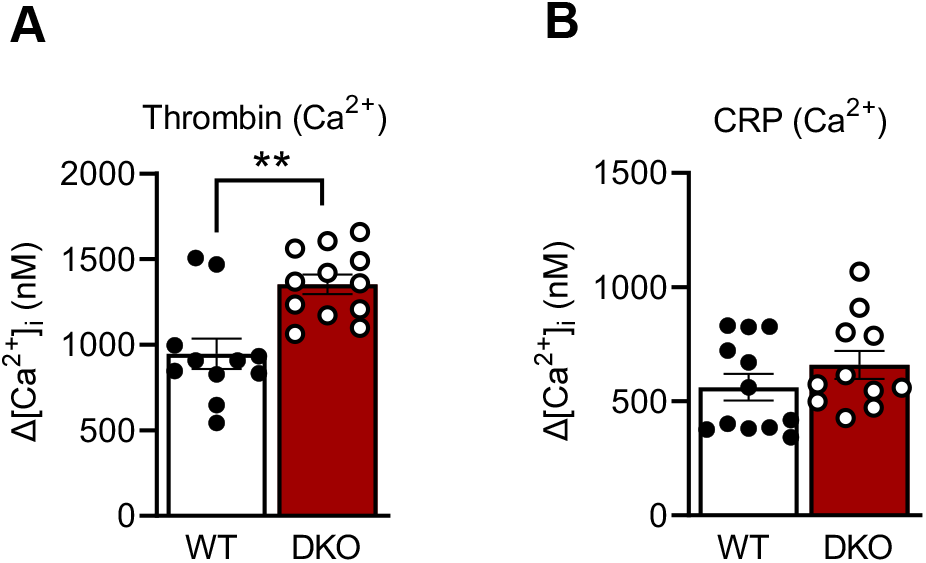
ZIP1/3 deficiency increases Ca^2+^ influx in response to thrombin but no effect in response to CRP. The increase in intracellular Ca^2+^ levels ([Ca^2+^]_i_) in washed platelets from WT and ZIP1/3 DKO mice stained with Fura-2 A/M after the stimulation with (A) Thrombin (0.01 U/ml) or (B) CRP (2 μg/ml) in the presence of 1 mM CaCl_2_; n=10-20 (Students t-test), ***P<0.001

**Figure 6:**
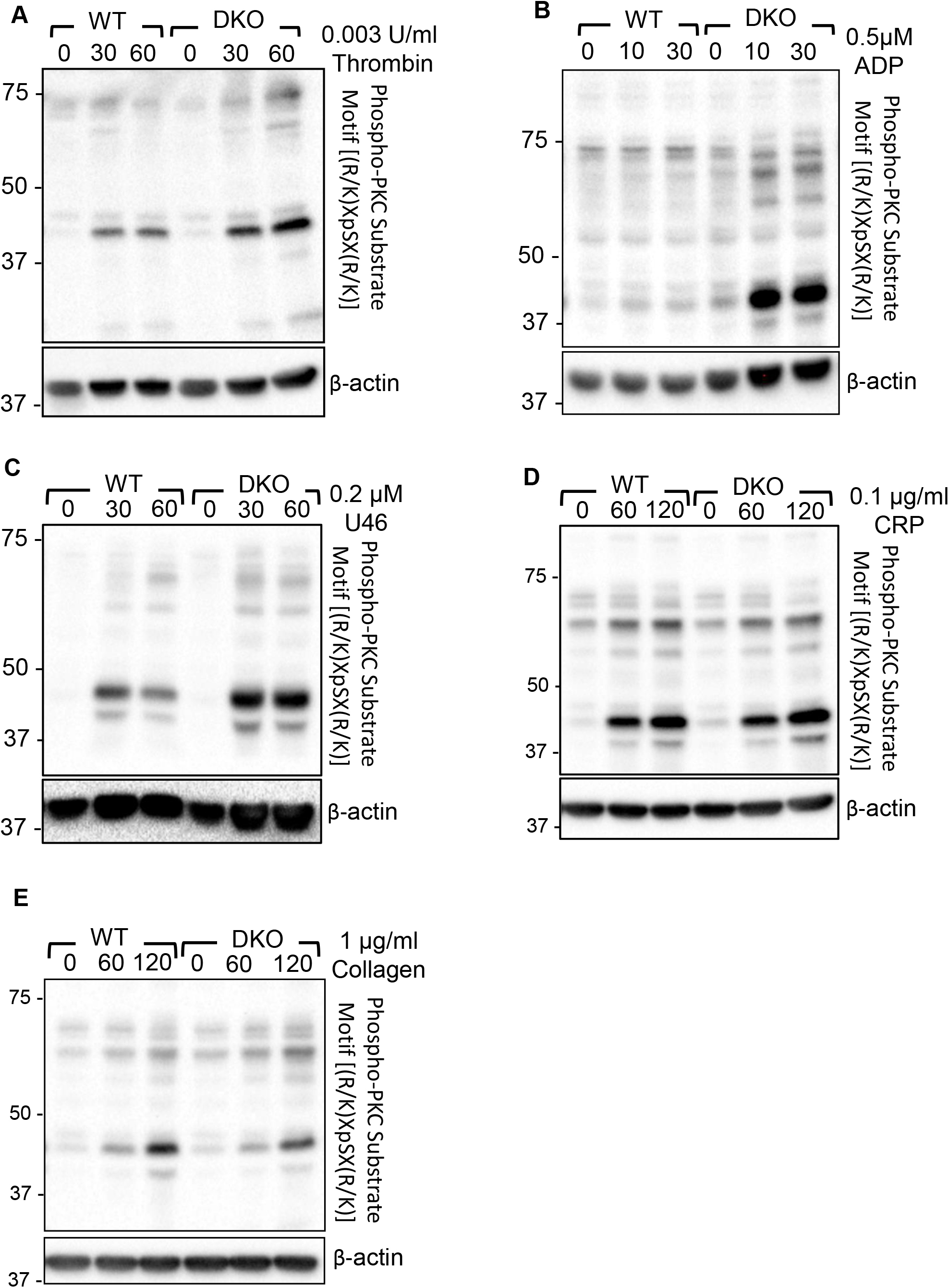
ZIP1/3 deficiency promotes PKC activation in murine platelets. Representative blots (n=3-5) show the levels of the phosphorylation of PKC substrates under resting (0s) conditions or upon stimulation with the GPCR agonists (A) Thrombin (0.003 U/ml for 30s and 60s); (B) ADP (0.5μM for 10s and 30s) or (C) U46619 (0.2μM for 30s and 60s), or the ITAM-coupled receptor agonists (D) CRP (0.1μg/ml for 60s and 120s) or (E) collagen (1μg/ml for 60s and 120s). Staining against β-actin served as loading control.

### Deficiency of ZIP1/3 transporters promotes thrombus formation

Zn^2+^ is considered to be an important contributor to haemostasis and thrombosis (17). According to this information and with respect to the observed hyperreactivity in aggregation (**Fig. 2A**), it was essential to determine the thrombotic potential *ex vivo* and *in vivo* in WT and ZIP1/3 DKO mice. In an *ex vivo* flow chamber assay, in which whole blood was perfused over collagen-coated coverslips at a shear rate of 1000s^−1^, we observed significantly bigger stable 3-dimensional platelet aggregates in case of blood from ZIP1/3 DKO mice, while surface coverage was only mildly affected (**Fig. 7A**). Consistent with these *ex vivo* findings, ZIP1/3 DKO mice formed clearly visible thrombi at earlier time points *in vivo* upon FeCl_3_-mediated injury of mesenteric arteries. The formed thrombus occluded mesenteric arteries of ZIP1/3 DKO mice faster as compared to wild-type mice confirming that platelets from ZIP1/3 DKO mice are hyperactive (**Fig. 7B**).

**Figure 7:**
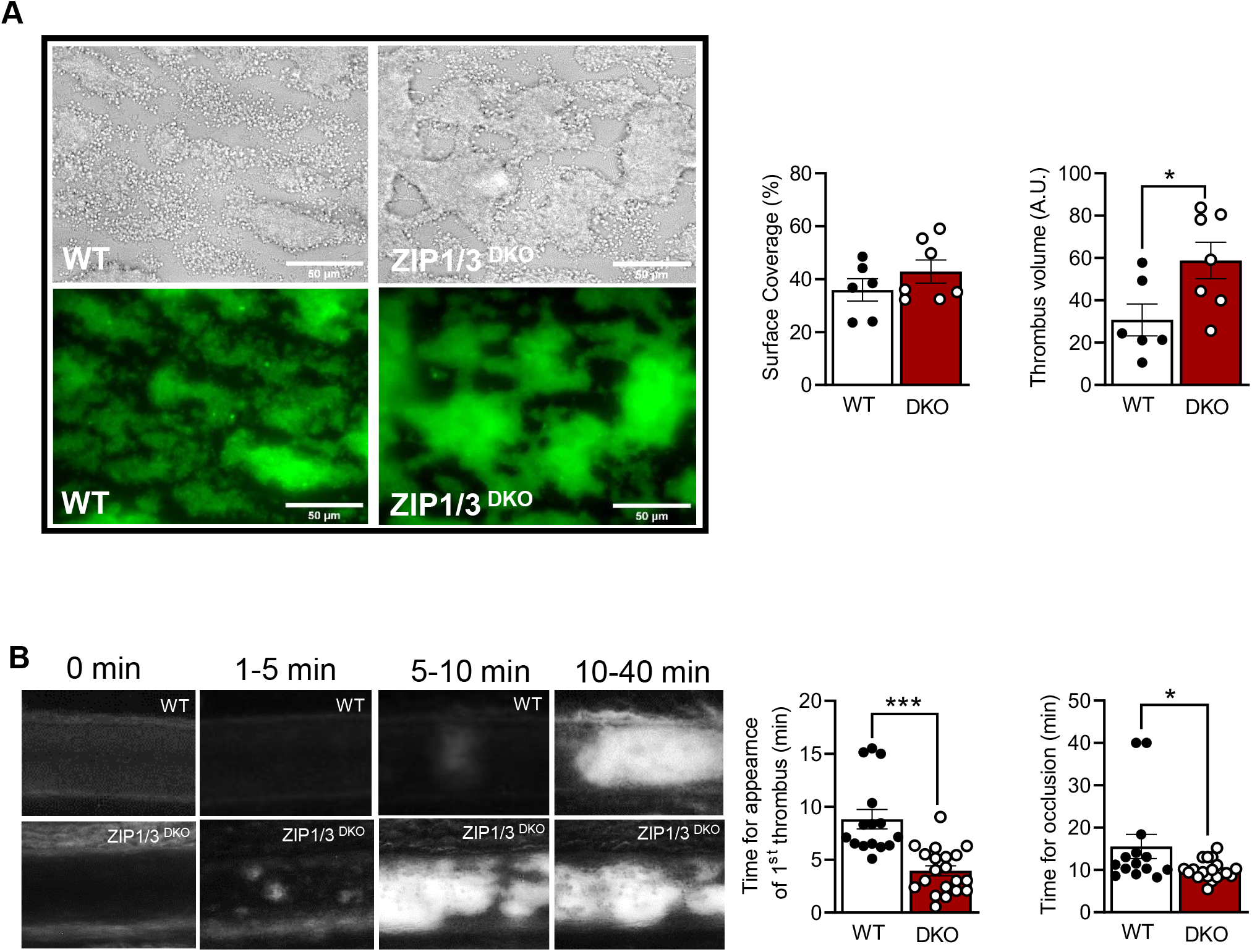
ZIP1/3 deficiency promotes *in vivo* and *in vitro* thrombus formation. (A) Adhesion and thrombus formation of platelets on collagen was assessed in a flow adhesion assay at a wall shear rate of 1000/s. Representative images and the quantification of the surface coverage and the relative thrombus volume are shown (n = 6; Students t-test; *P<0.05; Scale bar: 50 μm). (B) Representative images (left) and quantitative analysis (right) indicate the time for appearance of 1^st^ *in vivo* thrombus (> 10 μM) and time to occlusion at mesenteric arteries following induction of injury using FeCl_3_ (n=6-7 mice; each 2-3 arteries; Students t-test; *P<0.05, ***P<0.001).

## Discussion

Zn^2+^ homeostasis is mainly controlled through Zn^2+^ transporting proteins and metallothionein. Transporters of the ZIP superfamily are widely expressed in eukaryotic cells to import Zn^2+^ into the cytoplasm while ZnT proteins export Zn^2+^ out of the cytoplasm (31,32). The role of ZIP transporters in platelets, however, remained ill-defined. Interestingly, the global deletion of ZIP1 and ZIP3 provided the first direct proof that ZIP transporters are involved in the regulation of Zn^2+^ homeostasis in hippocampal neurons (40). The present study is, to our knowledge, the first to investigate the importance of ZIP transporters for platelet Zn^2+^ homeostasis and activation. Utilizing mice which globally lack ZIP1 and ZIP3, we provide evidence that these transporters selectively contribute to the control of platelet sensitivity towards GPCR agonists without affecting ITAM receptor agonists. Since ZIP1/3 DKO mice did not display alterations in platelet count, we conclude that megakaryopoiesis is independent of the presence of these transporters. Similarly, we did not observe any defects in other blood cells which is in line with the initial description of the mouse line (37). Due to the lack of specificity of all tested commercially available antibodies to detect ZIP1 and ZIP3, we confirmed the deletion of ZIP1 and ZIP3 by qPCR analysis in MKs generated *in vitro* from haematopoietic stem cells. We found that the deficiency of ZIP1 and ZIP3 altered Zn^2+^ homeostasis in platelets. While WT platelets release substantial amounts of free intracellular Zn^2+^ in response to thrombin, platelets from ZIP1/3 DKO mice showed a significantly delayed and less efficient release. As a net result the free [Zn^2+^]_i_ remained higher in thrombin-treated platelets from ZIP1/3 DKO mice. Potential mechanisms underlying the delayed and ineffective reduction of intracellular free Zn^2+^ in ZIP1/3 DKO platelets might involve a dysregulated expression or activity of ZnT export transporters or defective release of granules. A general defect in α-granule release, however, can be excluded since the exposure of P-selectin is even increased in response to thrombin in ZIP1/3-deficient platelets. Interestingly, recent work indicated that α-granules might not represent a homogeneous population of organelles, but rather comprise a group of subcellular compartments with unique composition and ultrastructure which might even release their content in an ordered manner (41–43). Whether ZIP1/3 may contribute to a selective release of platelet granules needs to be addressed in future experiments.

Not much is known about the downstream signalling mechanism by which ZIP1 and ZIP3 transporters control platelet activity. The hyper-responsiveness of platelet lacking ZIP1 and ZIP3 might be attributed to the increased retained pool of free intracellular Zn^2+^ which can act as a second messenger to enhance tyrosine phosphorylation events as shown in platelets (20), but also in nucleated cells, in particular mast cells (44), or might directly impact redox signalling (45). In addition, the enhanced Ca^2+^ entry in response to thrombin might likely affect a number or downstream signalling pathways. A common signalling molecule on which Zn^2+^ and Ca^2+^ pathways converge is PKC. It has been reported that Zn^2+^ augments PKC activity via increasing its association with regulatory ligand in lymphocytes and platelets (23). Besides Zn^2+^, Ca^2+^ activates PKC to regulate PS exposure in platelets(46). Clearly, there was an increased activation of PKC in ZIP1/3-deficient platelets in response to GPCR agonists, but not ITAM-coupled receptor agonists. According to these findings, there is a potential role of ZIP1 and ZIP3 in orchestrating GPCR signalling pathways in platelets. Taken in consideration that PKC is essential for thrombus formation (47), PKC activation could explain the observed promoted thrombus formation in ZIP1/3 DKO mice. Given that PKC activation also mediates platelet granule secretion (47,48), the data are consistent with the selective increase of P-selectin in platelets from ZIP1/3 DKO mice in response to GPCR agonists, particularly thrombin. Besides granule secretion, PKC activation is a positive regulator of integrin activation (49). Indeed, flow cytometric analyses confirmed an increased activation of α_IIb_β_3_ in response to thrombin stimulation in ZIP1/3 DKO platelets which is in line with its role to promote platelet aggregation.

In aggregometry experiments as well as in flow cytometry experiments of α_IIb_β_3_ integrin activation and degranulation, ZIP1/3-deficient platelets were hyperresponsive in response to threshold concentrations of GPCR agonists, but not towards agonists of the ITAM-coupled receptor GPVI, namely CRP and collagen. Nevertheless, platelets from ZIP1/3 DKO formed larger thrombi when perfused over a collagen-coated surface. This is most likely a consequence of second wave mediators, like TxA2 and ADP, which are generated and released upon platelet activation and amplify platelet activation via GPCR signalling in order to promote thrombus growth. Indeed, it has been shown that co-infusion of ADP and the stable TxA2 analogue U46619 into the flow chamber enhances thrombus formation (50), hence arguing that the observed phenotype is due to enhanced GPCR signalling of ZIP1/3 DKO platelets. Of note, thrombin does not contribute to this effect in this experimental setup since the presence of heparin in the flow chamber abolishes its activity. Alternatively, the prevailing shear forces and the presence of plasma factors might also contribute to the experimental outcomes. GPCR-mediated signalling pathways are also prevailing in our model of FeCl_3_-induced vessel injury (51–53), which is triggered by oxidative damage of vessel wall and blood cells and flocculation of blood proteins, while there is only limited, if any, collagen exposure inside the vessel (52,54,55). Therefore, the enhanced thrombus formation in ZIP1/3 DKO mice is again in line with the hyperresponsiveness of their platelets towards GPCR agonists. Noteworthy, since a global KO model was used, we cannot exclude an impact of other cell types, e.g. endothelial cells and blood cells, as well as systemic alterations on the experimental outcome. An additional aspect that might also be of relevance is the role of Zn^2+^ in altering fibrin network structure and its mechanical properties (56). To what extent the altered Zn^2+^ release of ZIP1/3 DKO influences thrombus formation and stabilization via modification of the fibrin network in this assay warrants further investigation.

Taken together, our study for the first time attributes an important regulatory role to zinc transporters in platelet activation. *In vivo*, ZIP1/3 DKO mice develop faster occlusive thrombi in response to vascular injury. *In vitro* data show that platelets from ZIP1/3-deficient mice are hyperreactive to threshold concentrations of GPCR agonists, in particular thrombin, while responses to ITAM-coupled receptor agonists appear to be unaffected. Future studies are aimed at the understanding of the molecular mechanisms which determine this unexpected exclusiveness.

## Author contributions

AE, PÖ, ME, FK, CK, UG, TV and HMH performed experiments and analysed data. BN discussed results and provided scientific input and technical equipment throughout the study. MB generated the mice. AE, TV, HMH wrote the manuscript with input from all the authors. AE and HMH conceived the study and designed research.

## Acknowledgements

This work was supported by the Deutsche Forschungsgemeinschaft (DFG, German Research Foundation) (project number 374031971-TRR 240/project A09). We thank Donata Dorbath, Sandra Umbenhauer and Birgit Midloch for excellent technical assistance.

