## Supplementary Figure 1 for "Loss of zinc transporters ZIP1 and ZIP3 augments platelet reactivity in response to G protein-coupled receptor agonists and accelerates thrombus formation *in vivo*"

### Supplementary Figure 1 (Elgheznawy *et al.*)

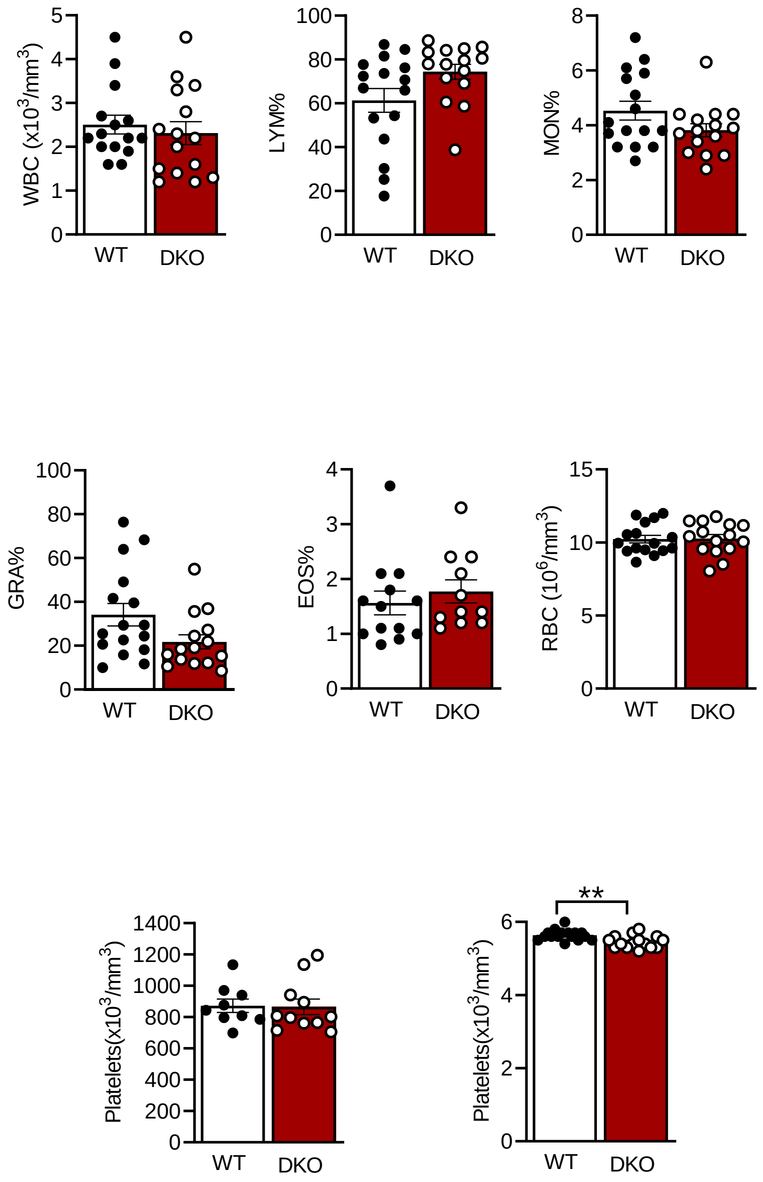


**Supplementary figure 1: ZIP1/3 deficiency has no effect on whole blood count**. Analyses of blood parameters were performed using a scilVetabcPlus+ hemacytometer from EDTA blood (scilVet, scil animal care company GmbH, Viernheim, Germany).
