## Supplementary Figure 2 for "Loss of zinc transporters ZIP1 and ZIP3 augments platelet reactivity in response to G protein-coupled receptor agonists and accelerates thrombus formation *in vivo*"

### Supplementary Figure 2 (Elgheznawy *et al.*)

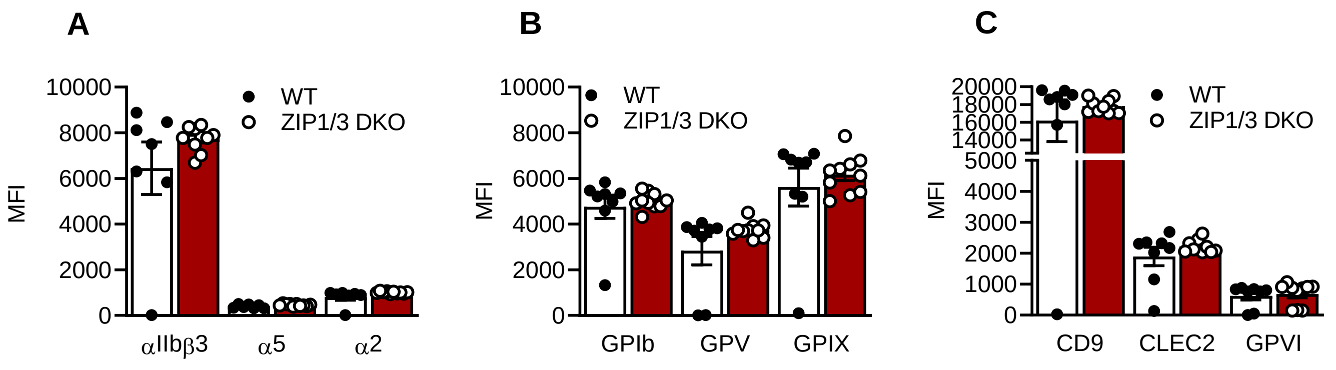


**Supplementary Figure 2: ZIP1/3 deficiency does not affect platelet glycoprotein or integrin cell surface levels.** Flow cytometric analysis of the cell surface expression of the indicated integrins (A), the GPIb-V-IX complex (B) and CD9, CLEC2, GPVI using directly fluorescently labelled specific antibodies in saturating amounts.
